# Developing peptide-based fusion inhibitors as an antiviral strategy utilizing Coronin 1 as a template

**DOI:** 10.1101/2024.07.04.602150

**Authors:** Manbit Subhadarsi Panda, Bushra Qazi, Vaishali Vishwakarma, Gourab Prasad Pattnaik, Sourav Haldar, Hirak Chakraborty

## Abstract

Enveloped viruses can enter the host cells by endocytosis and subsequently fuse with the endosomal membranes, or fuse with the plasma membrane at the cell surface. The crucial stage of viral infection, regardless of the route taken to enter the host cell, is membrane fusion. The present work aims to develop a peptide-based fusion inhibitor that prevents membrane fusion by modifying the properties of the participating membranes, without targeting a protein. This would allow us to develop a fusion inhibitor that might work against a larger spectrum of enveloped viruses as it does not target any specific viral fusion protein. With this goal, we have designed a novel peptide by modifying a native sequence derived from coronin 1, a phagosomal protein, that helps to avoid lysosomal degradation of mycobacterium-loaded phagosomes. The designed peptide, mTG-23, inhibits ∼ 30-40% fusion between small unilamellar vesicles containing varying amounts of cholesterol by modulating the biophysical properties of the participating bilayers. As a proof of principle, we have further demonstrated that the mTG-23 inhibits Influenza A virus infection in A549 and MDCK cells (with∼ EC_50_ of 20.45 µM and 21.45 µM, respectively), where viral envelope and endosomal membrane fusion is a crucial step. Through a gamut of biophysical and biochemical methods, we surmise that mTG-23 inhibits viral infection by inhibiting viral envelope and endosomal membrane fusion. We envisage that the proposed antiviral strategy can be extended to other viruses that employ a similar modus operandi, providing a novel pan-antiviral approach.

## Introduction

Membrane fusion plays vital roles in different cellular episodes like fertilization, neurotransmission, cellular trafficking, cell division, exo- and endocytosis, and entry of enveloped viruses.^1–5^ It is a process in which two distinct lipid bilayers merge their hydrophobic cores resulting in the formation of an interconnected structure.^6^ Enveloped viruses utilize the membrane fusion process to transfer their genome inside the host cells and multiply using the host replication machinery.^1, 7^ A special class of viral surface glycoproteins in the viral envelope play a crucial role in binding of the virus at the host cell membrane and inducing fusion. A recent review summarized the efforts made to block the entry of enveloped viruses into the host cell by targeting the viral fusion proteins and cell surface receptors.^8^ Quite a few studies demonstrated that short peptides mimicking the sequences of either the N-heptad or C-heptad region of the fusion protein inhibit the six-helix bundle formation by interacting with the complementary heptad regions of the fusion protein and block fusion between the virus and host cells.^9–11^ Moreover, inhibitors have been designed to directly block the interaction between the receptor binding domain of viral fusion protein and cellular receptors or by mimicking the cellular receptors so that the virus will attach itself to the inhibitor peptide instead of cellular receptors.^12–14^ Some of the existing HIV fusion inhibitors that target gp140-CD4 interactions are PRO 2000, and B69 (3HP-β-LG).^15, 16^ There exist numerous inhibitor peptides that are effective for various deadly viruses like HIV, the most common clinical peptide-based drug that was discovered was Enfuvirtide (T20).^17^ A peptide derived from the HR2 (membrane proximal) region of the spike protein of severe acute respiratory syndrome coronavirus (SARS-CoV) interacts with the HR1 (membrane distal) region and displays an inhibitory effect.^18^ Inhibitors have been further designed to block a serine protease, TMPRSS2, that activates the fusion domain of the SARS-CoV spike protein by inducing successful cleavage between receptor binding and the fusion domain of the fusion protein.^19^ However, these inhibitors are limited by their applications against a particular virus as they target the fusion proteins or cell surface receptors, which vary across viruses. The traditional approach of the ‘one bug-one drug’ in antiviral therapy might be inadequate to combat the continuously emerging and reemerging viral diseases.^20, 21^ However, the broad-spectrum entry inhibitors cannot be designed by targeting the viral fusion proteins because of their variability in viruses. Nonetheless, the fusion process occurs through some common pathways in terms of the evolution of membrane structure and conformation.^2^ Small molecules or peptides can successfully hinder the conformational evolution of membranes and inhibit the fusion.^21, 22^

In general, intercellular pathogens like *mycobacterium* recruit a tryptophan-aspartic acid coat protein, coronin 1, to escape lysosomal degradation by preventing endosome-lysosome fusion.^23^ Interestingly, coronin 1 contains tryptophan-aspartic acid (WD) repeats, and they are conserved across species.^24^ Additionally, it controls several F-actin-dependent activities, including micropinocytosis, phagocytosis, and motility.^25^ We have earlier shown that a coronin 1-derived WD-containing peptide, TG-23 (TT**WD**SGFCAVNPKFVALICEASG), inhibits polyethylene glycol (PEG)-induced fusion between small unilamellar vesicles (SUVs).^22^ The peptide inhibited pore opening by stabilizing the hemifusion intermediate by reducing water percolation at the acyl chain region. However, the ability to reduce water percolation disappears with increasing concentration of membrane cholesterol so as its inhibitory efficiency.^26, 27^

In the present work, we designed a new WD-containing peptide, mTG23, by modifying the sequence of TG-23, which demonstrates fusion inhibitory ability irrespective of the cholesterol content of the membrane as opposed to what we observed in the case of TG-23. As TG-23 contains a WD repeat near the N-terminal region and demonstrates inhibitory efficacy in the membrane devoid of cholesterol,^22^ we modified the TG-23 peptide by introducing another WD repeat near the C-terminal region, (TT**WD**SGFCAVNPKFVALIC**DW**SG), and evaluated its fusion inhibitory efficacy in PEG-mediated fusion of SUVs. We have measured the peptide-induced changes in the organization and dynamics at the interfacial region of the membrane by utilizing steady-state and time-resolved fluorescence methods. To evaluate whether mTG-23’s fusion inhibitory effect translates to antiviral activity, we have tested its *in vitro* efficacy against the Influenza A virus. Our results demonstrate that mTG-23 inhibits pore opening in the PEG-induced fusion of SUVs independent of membrane cholesterol level and prevents Influenza A Virus infection in human lung adenocarcinoma epithelial (A549) and Madin-Darby canine kidney (MDCK) cells.

## MATERIALS AND METHODS

### Materials

1,2-dioleoyl-sn-glycero-3-phosphocholine (DOPC), 1,2-dioleoyl-sn-glycero-3-phosphoethanolamine (DOPE), and cholesterol (CH) were procured from Avanti Polar Lipids (Alabaster, AL). N-(7-nitrobenz-2-oxa-1,3-diazol-4-yl)-1,2-dihexadecanoyl-sn-glycero-3-phosphoethanolamine (NBD-PE), and Lissamine rhodamine B 1,2-dihexadecanoyl-sn-glycero-3-phosphoethanolamine (Rh-PE) were purchased from Thermo Fisher Scientific (Waltham, MA). Triton X-100 (TX-100), poly (ethylene glycol) of molecular weight 7000-9000 gm (PEG 8000), sephadex G-75, dipicolinic acid (DPA), and trimethyl ammonium derivative of 1,6-diphenyl-1,3,5-hexatriene (TMA-DPH) were procured from Sigma Aldrich (St. Louis, MO). N-[Tris(hydroxymethyl)-methyl]2-2-aminoethane sulfonic acid (TES) and terbium chloride were obtained from Alfa Aesar (Haverhill, MA). Calcium chloride and ethylenediaminetetraacetic acid (EDTA) were obtained from Merck (India), whereas sodium chloride was obtained from Fisher Scientific (India). UV-grade solvents such as chloroform and methanol have been procured from Spectrochem (India). All the chemicals used in the work are of 99% purity. All chemicals utilized in this work have more than 99% purity. Water was purified through an Adrona Crystal (Latvia) water purification system.

### Peptide Synthesis

The mutant TG-23 peptide sequence is TT**WD**SGFCAVNPKFVALIC**DW**SG which was chemically synthesized and purified by Biochain Incorporated (New Delhi, India). The peptides were synthesized by the solid phase approach using Fmoc chemistry, which is discussed elsewhere.^28, 29^ Mass spectrometry has been utilized to characterize the peptide (**Supp. Fig. 1**) that was isolated via HPLC (**Supp. Fig. 2**). Small aliquots of the peptide stock solutions were added to the vesicle suspensions after they were prepared in DMSO. The DMSO concentration was always less than 0.5 % (v/v), and it had no discernible impact on membrane shape or fusion.^30^

### Preparation of Vesicles

Using the sonication method, small unilamellar vesicles (SUVs) were prepared from either DOPC/DOPE/DOPG (60/30/10 mol %), DOPC/DOPE/DOPG/CH (50/30/10/10 mol %), DOPC/DOPE/DOPC/CH (40/30/10/20 mol %).^31^ Lipids at this appropriate molar ratio, in chloroform were dried overnight in a vacuum desiccator. For uniform dispersion of lipids, the dried films were hydrated and vortexed in the assay buffer for 1 h. The column (Sephadex G-75) and experimental buffer were comprised of 10 mM TES, 100 mM NaCl, 1 mM EDTA, and 1 mM CaCl_2_ at pH 7.4. Then to prepare SUVs, the hydrated lipids were sonicated using a Dr. Hielscher model UP100H (Germany) probe sonicator, as documented previously.^22^ Sonicated samples were centrifuged at 15,300 g for 10 minutes to remove aggregated lipids and titanium particles. All lipid mixing, content mixing, and leakage experiments were performed with 200 μM lipid. Using dynamic light scattering (Malvern Zetasizer Nano ZS-90), the average hydrodynamic radii of the vesicles were determined to be 50-60 nm with a polydispersity index of less than 0.2 (data not shown). Small aliquots of the probe were added from their respective stock solutions prepared in DMSO into working solutions. The DMSO content was always less than 1% (v/v), and this small quantity of DMSO was established not to produce any detectable effect on membrane structure.^30^

### Steady-State Fluorescence Anisotropy Measurements

Steady-state fluorescence anisotropy measurements were performed using a Hitachi F-7000 (Japan) spectrofluorometer with a 1cm path-length quartz cuvette. TMA-DPH was excited at 360 nm, and its emissions were monitored at 428 nm. In all measurements excitation and emission slits with a nominal bandpass of 5 nm were used. All measurements were conducted in a buffer containing 10 mM TES, 100 mM NaCl, 1 mM CaCl_2_, and 1 mM EDTA, pH 7.4 at 37 °C. Anisotropy values were calculated using the following equation^32^

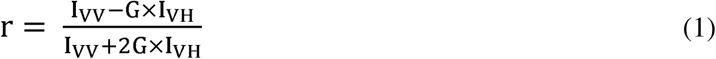

where, G = I_HV_/I_HH_, (grating correction or G-factor), I_VV_ and I_VH_ are the measured fluorescence intensities with the excitation polarizer vertically oriented and the emission polarizer vertically and horizontally oriented, respectively.

### Lipid Mixing Assay

The transfer of lipid (Lipid Mixing) during PEG-mediated vesicle fusion was examined based on the change in FRET efficiency between FRET lipid pairs: NBD-PE (donor) and Rh-PE (acceptor).^22, 33^ FRET dilution as a function of time is reflected as a marker to measure the kinetics of lipid transfer (mixing) between two vesicles.^22^ We prepared a set of vesicles comprising FRET lipid pairs in equal concentration (0.8 mol %) and indicating maximum FRET. The probe-containing vesicles were mixed with probe-free vesicles at a ratio of 1:9.^34^ 6% (w/v) PEG was added to 200 μM lipid vesicles to initiate the lipid mixing process, which was measured by observing the reduction in FRET efficiency via a rise in donor intensity. To evaluate the effect of inhibitor peptide, 2 μM peptide was added to the mixture of probe-containing and probe-free (1:9) vesicles 10 mins before the addition of PEG to monitor the lipid mixing. The emission intensity of the donor (NBD-PE) was examined with Hitachi F-7000 (Japan) spectrofluorometer, keeping the excitation and emission wavelengths fixed at 460 nm and 530 nm, respectively. Throughout the experiment, a minimum slit of 5 nm was used on both the excitation and emission sides. Each experiment was repeated at least three times. The percentage of lipid mixing was calculated using the following equation^33^

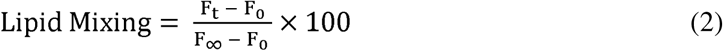

where ‘F_0_’, ‘F_t_’, and F∞’ are the fluorescence intensities at the zeroth time, time = t and time = ∞, respectively. F∞ has been measured in the presence of TX-100, which is considered the complete mixing of lipids.

### Content Mixing Assay

We have monitored the content mixing as proposed by Wilschut et al. using Tb^3+^ and DPA.^35,36^ Vesicles were prepared either in 80 mM DPA or 8 mM TbCl_3_. The untrapped DPA and TbCl_3_ were removed from the external buffer of the vesicles using a Sephadex G-75 column equilibrated with assay buffer (10 mM TES, 100 mM NaCl, 1 mM EDTA,1 mM CaCl_2_ at pH 7.4). 6% (w/v) PEG was added to 200 μM lipid vesicles to induce content mixing. 2 μM peptide was added to a 200 μM mixture (1:1) of Tb^3+^ and DPA-containing vesicles 10 mins before the addition of PEG to monitor the content mixing. The content mixing was measured in terms of an increase in fluorescence intensity due to the formation of the Tb/DPA complex with time. The excitation and the emission wavelength for the content mixing assay were used as 278 nm and 490 nm, respectively. A minimum slit width of 5 nm in both the excitation and emission side was used for all measurements. Each experiment was repeated at least three times. The percentage of content mixing was calculated in the following way:

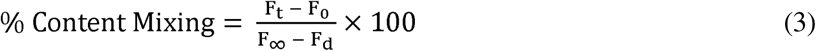

where ‘F_0_’ and ‘F_t_’ are the fluorescence intensities at zeroth time and time = t, respectively. ‘F∞’ and ‘F_d_’ have been calculated from the fluorescence intensity of the leakage sample at a zeroth time (which gives the highest intensity as all Tb are being complexed with DPA) and in the presence of detergent, respectively. ‘F∞’ implies the highest fluorescent signal when Tb^3+^ and DPA are in the same vesicular compartment and all Tb^3+^ are in a complex form with DPA. Upon detergent addition, Tb^3+^ and DPA will come out from the vesicular compartment and be diluted in the bulk solution, resulting in dissociation of the complex and minimum fluorescence intensity (F_d_).

### Leakage Assay

The leakage assay was carried out by monitoring the decrease in fluorescence intensity of the vesicles containing both TbCl_3_ and DPA in the presence of PEG or PEG and peptide.^35, 36^ 8 mM TbCl_3_ prepared in 10 mM TES and 100 mM NaCl, pH 7.4, and 80 mM DPA which is prepared in 10 mM TES, pH 7.4, were co-encapsulated in the vesicles. The external TbCl_3_ and DPA were removed using a Sephadex G-75 column equilibrated with assay buffer (10 mM TES, 100 mM NaCl, 1 mM EDTA, 1 mM CaCl_2_ at pH 7.4).^34^ 6% (w/v) PEG was added to 200 μM lipid Tb^3+^ and DPA-containing vesicles to induce content leakage. To evaluate the effect of inhibitor peptide, 2 μM peptide was added to the vesicles 10 mins before the addition of PEG to monitor the content leakage. The maximum leakage (100%) was characterized by the fluorescence intensity of a co-encapsulated TbCl_3_/DPA vesicle treated with 0.1% (w/v) Triton X-100. The excitation and the emission wavelength were fixed at 278 nm and 490 nm, respectively. Slits of 5 nm were used in both the excitation and emission sides throughout the experiment. Each experiment was repeated at least three times. The percentage of content leakage was calculated in the following way:

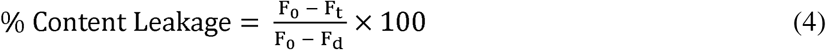

where ‘F_0_’ and ‘F_t_’ and ‘F_d_’ are the fluorescence intensities at the zeroth time, time = t, and in the presence of detergent, respectively.

### Time-resolved Fluorescence Measurements

Fluorescence lifetimes were measured from time-resolved fluorescence intensity decays using the IBH 5000F Nano LED equipment (Horiba, Edison, NJ) with Data Station software in time-correlated single photon counting (TCSPC) mode, as mentioned earlier.^22, 30^ A pulsed light-emitting diode (LED) was applied as the excitation source. The LED generated an optical pulse at 340 nm (for exciting TMA-DPH), with a pulse duration of less than 1.0 ns, and was run at a rate of 1 MHz. The Instrument Response Function (IRF) was measured at the respective excitation wavelength using Ludox (colloidal silica) as a scatterer. Ten thousand photon counts were collected in the peak channel to improve the signal-to-noise ratio. All experiments were performed using emission slits with an 8-nm bandpass for TMA-DPH. Data were stored and analyzed using DAS 6.2 software (Horiba, Edison, NJ). Fluorescence intensity decay curves were deconvoluted with the instrument response function and analyzed as a sum of exponential terms as follows:

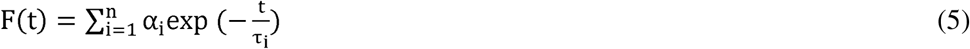

A considerable plot was found with a random deviation around zero with a maximum χ^2^ value of 1.2 or less. Intensity-averaged mean lifetimes τ_avg_ for n-exponential decays of fluorescence were calculated from the decay times and pre-exponential factors using the following equation^32^

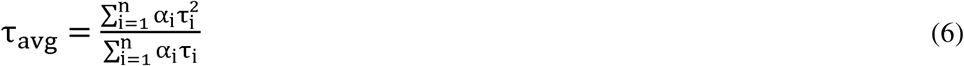

where α_i_ is the fraction that shows lifetime τ_i_.

### Calculation of Apparent Rotational Correlation Time

Apparent rotational correlation time delivers information about the ease of rotational motion of fluorophores during their lifetime, which is affected by the rigidity of its immediate microenvironment. Therefore, we can acquire information about membrane viscosity in the interfacial region by monitoring the apparent rotational correlation time (θ_c_) of TMA-DPH from steady-state anisotropy and the average lifetime values using Perrin’s Equation^32^ as follows:

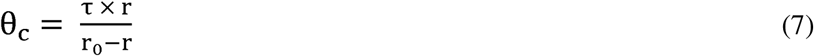

where ‘τ’ is the average lifetime of the probe and r is anisotropy. The constant r_0_ value was considered as 0.4 for TMA-DPH.^32^

### Cell lines, Viruses, and Antibodies

MDCK and A549 cells were obtained from ATCC. MDCK and A549 cells were maintained in DMEM (GIBCO) with 10 % FBS (HIMEDIA) containing 100 μg/ml penicillin (HIMEDIA) and 100 μg/ml streptomycin (HIMEDIA). Cells were grown in a humidified atmosphere (95% humidity) at 37 °C and 5% CO_2_. Trypsin was obtained from HIMEDIA. Influenza A virus (strain X-31, A/Aichi/68, H3N2) was a kind gift from Dr. Joshua Zimmerberg at NICHD/ NIH, Bethesda. A mouse monoclonal Anti M1 antibody (ab22396) and a rabbit anti-mouse IgG H&L [HRP] (ab6728) were obtained from Abcam. An HRP-conjugated anti β actin (A3854) was obtained from Sigma.

### Viral Infectivity and Peptide Efficacy Assay

X-31 (H3N2) Influenza viral infection was carried out in A549 and MDCK cell lines. The *in vitro* antiviral effect of mTG-23 was evaluated by quantifying the reduction in cytopathic effect induced by the Influenza A virus in A549 and MDCK cells. 1.5 X 10^4^ (A549 or MDCK) cells were seeded in each well of a 96-well tissue culture plate and were allowed to grow as a monolayer for 24 hours. After adherence, the cells were incubated with different concentrations of the peptide for 48 hours at 37 °C and 5% CO_2_. After 48 hours cell viability was assessed by MTT assay. For infection in the presence of the peptide, mTG-23 was two-fold serially diluted (from a 1 mM stock in DMSO) in serum-free media (SFM) containing trypsin (2.5 µg/ml). Equal amounts of Influenza A virus from stock were pre-mixed with different amounts of the peptide in SFM. A549 and MDCK cells were then infected with peptide-virus mixtures for 45 minutes at 37°C, 5% CO_2._ After that, infection media was removed followed by washing the cells with PBS to remove unbound virus and then supplemented with growth media containing 0.1% FBS, 0.3% BSA, and trypsin (2.5 µg/ml) and incubated for 48 hours at 37°C, 5% CO_2_. After 48 hours, 10μl of MTT (SRL) (from a stock of 5 mg/ ml in PBS) was added to each well and the plate was incubated for 4 hours at 37°C, 5% CO_2._ After that 100 μl of DMSO was added in each well to dissolve the formazan crystals produced in the process and absorbance was measured at 570 nm. The wells containing only cells and cells infected with the virus were considered as cell control and virus control, respectively. Cell viability was determined by considering the cell control absorbance corresponds to 100 % viability. The percentage of cytopathic effect (CPE) inhibition was defined as

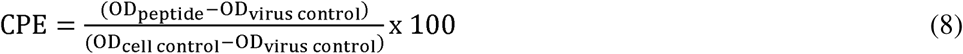

where OD_peptide_ is the absorbance in the presence of virus and peptide, OD_virus control_ is the absorbance in the presence of virus only and OD_cell control_ is the absorbance of cell control wells.^37^ Each O.D. value was background subtracted. Reduction in CPE is plotted as a function of peptide concentration and fitted to a dose-response model in GraphPad Prism 6 (San Diego, CA, USA), and 50% effective concentration (EC_50_) was extracted (see **Fig. 5**). The highest amount of DMSO used was 5 % (v/v) and showed no significant effect.

### Western blot analysis

To determine the level of viral protein expression, MDCK and A549 cells were seeded in a 6-well plate at 0.25×10^6^ cells / well and infected with Influenza A virus in the absence and presence of varying amounts of mTG-23 as described above. After 48 hours of infection, cells were observed for cytopathic effect (See **Fig. 6 A**). Media was harvested for HA titer determination. After removing the media, cells were washed with ice-cold PBS and scrapped in a lysis buffer (RIPA-HIMEDIA: 50 mM Tris-HCl pH 7.4, 1% Triton X-100, 1% sodium deoxycholate, 1 mM EDTA, 0.1% SDS) in the presence of a protease inhibitor (SIGMA). To prepare cell lysate cells were subjected to sonication for 90 seconds (pulse 15, Amp1 60%) (Fisher Scientific) and then pelleted by centrifugation at 16000 RCF for 20 minutes at 4 °C. The supernatant was collected and stored at -20 °C. Total protein concentration was determined by a bicinchoninic acid assay using a kit (Thermofisher, USA). Protein lysates were separated on an SDS-PAGE (12%) and then transferred to a PVDF membrane (Bio-Rad). The membrane was blocked by dipping in 3% skimmed milk (Bio-Rad) in PBS containing 0.1% Tween 20 (Amresco) (1xPBST) for 1.5 h with slow rocking at room temperature. IAV-associated matrix protein (M1) was detected by an anti-M1 antibody (1:1000) as primary and an HRP conjugated rabbit anti-mouse IgG H&L (1:10,000) was used as a secondary antibody. The same blots were used after stripping for estimating β-actin as a loading control using an HRP-conjugated primary antibody (1:10,000).

### Hemagglutination Assay

Hemagglutination assay (HA) is a quick way to measure the amount of virus present, given as the HA titer. This test relies on the ability of Influenza Virus Hemagglutinin to clump together and prevent red blood cells (RBC) from precipitation. In the absence of virus particles, RBCs settle at the bottom of the well, forming a red dot (refer to **Fig. 5B**). However, when virus particles are present, the RBCs form a lattice (clump together) due to the interaction between the HA proteins of the virus particles and the RBCs. This prevents the RBCs from precipitating, resulting in the absence of a red dot. The amount of virus present is linked to the extent of lattice formation (agglutination), as indicated by the dilution factor.

This factor represents the number of the last well (in the 2-fold serial dilution) that shows agglutination (absence of a red dot). Beyond a certain dilution fold, a red dot appears depending on the amount of virus present in the sample. A higher titer indicates a higher amount of virus. The assay was conducted using chicken RBC according to the reported procedures.^38, 39^ Briefly, RBCs were isolated by diluting blood in PBS and then centrifuging at 2500 RCF for 10 minutes at 25°C, and this process was repeated 3-4 times until a clear supernatant was obtained. For the HA assay, 50µl of 1X PBS was added from the 2nd to 12th well of a 96-well U-bottom plate. Then, 100µl of the infected media (with and without the peptide) was added to the first well and 2-fold serially diluted across the row. 50µl of infected media was removed from the last well. Subsequently, 50µl of standardized RBC (5% RBC in PBS) was added to each well and incubated for 30 minutes at room temperature. The last well which showed agglutination (absence of a red dot) was observed and recorded. The titer (dilution factor) of the virus control is then compared with that in the presence of different concentrations of the peptide, as depicted in **Fig. 5B**.

## Results

### Partitioning of mTG-23 peptide in membranes

The partitioning of mTG-23 in the membranes was evaluated by measuring the fluorescence spectra of tryptophan present in the mTG-23 peptide in the buffer and the presence of membranes of varying lipid compositions. The significant increase in fluorescence intensity implies the partitioning of the peptide in membranes. The fluorescence intensity does not depend on the lipid composition, which suggests a similar partition coefficient (or binding K_d_) of the peptide in the membranes having different cholesterol concentrations. Therefore, the change in the ability of the peptide to modulate membrane fusion is not due to its variable partitioning in the membrane rather it is due to its ability to alter the organization and dynamics of membranes with different cholesterol concentrations.

### Effect of mTG-23 peptide on PEG-mediated vesicle fusion

We have assessed the inhibitory effect of mTG-23 peptide in PEG-mediated fusion of SUVs. The fusion between SUVs was induced by adding 6% (w/v) PEG. It is known that 6% (w/v) PEG does not affect the physical properties and depth of penetration of the peptide to the membrane.^22, 29, 40^ Using the methodologies described in the materials and methods section, fusion observables like lipid mixing (LM), content mixing (CM), and leakage (L), were measured during the fusion of SUVs with lipid compositions of DOPC/DOPE/DOPG (60/30/10 mol%), DOPC/DOPE/DOPG/CH (50/30/10/10 mol%) and DOPC/DOPE/DOPG/CH (40/30/10/20 mol%). PEG-induced vesicular fusion in the absence of peptide in a membrane has been considered as the control for that membrane system, and the inhibitory efficiency of the peptide was calculated against the corresponding control. The inhibitory efficiency of the peptide was measured in PEG-mediated fusion by keeping the lipid-to-peptide concentration ratio of 100:1 to maintain the physiological relevance of low concentration of peptide.

**Fig. 2(A-C)** shows the time courses of LM, CM, and L respectively in the absence and presence of mTG-23 peptide during the fusion of SUVs having lipid composition DOPC/DOPE/DOPG (60/30/10 mol%). The percentages of LM, CM, and L have been calculated using Eqns. (2), (3), and (4), respectively described in the materials and method section. The extent of LM, CM, and L at infinite time has been calculated by fitting the kinetic data in a double exponential equation using the SigmaPlot (San Jose, CA) software package and the results have been shown in Table 1. Interestingly, mTG-23 promotes LM (by around 20%), however, inhibits CM and L by approximately 40% and 25%. This indicates that the peptide is stabilizing the hemifusion intermediate and reduces the ability of pore opening. Generally, a fusogenic system demonstrates a higher extent of L, therefore reduction of L in the presence of mTG-23 further supports the inhibition of content mixing result. Similar experiments were carried out in DOPC/DOPE/DOPG/CH (50/30/10/10 mol%) and DOPC/DOPE/DOPG/CH (40/30/10/20 mol%) membranes in the absence and presence of mTG-23 peptide. **Fig. 2(D-F)** demonstrates the kinetics of LM, CM, and L in the absence and presence of mTG-23 peptide in DOPC/DOPE/DOPG/CH (50/30/10/10 mol%) membranes. The peptide does not have much impact on LM (increases by 6%), however, displays inhibition of CM (by 30%) and L (by 40%). A comparable trend in LM (enhanced by 8%) and CM (inhibited by 33%) was observed in the presence of the peptide in DOPC/DOPE/DOPG/CH (40/30/10/20 mol%) membranes (Fig. 2**(G-I)**). Taken together, our results demonstrate that the mTG-23 peptide partially promotes LM vis-à-vis hemifusion formation but inhibits pore opening irrespective of the concentration of membrane cholesterol. This result is contrary to our observation of the TG-23 peptide, which inhibited pore opening in the absence of cholesterol, however, loses its inhibitory efficiency with increasing concentration of membrane cholesterol.^26, 27^ Therefore, the C-terminus mutated mTG-23 peptide can be considered a better candidate as an entry inhibitor against a larger spectrum of lipid compositions. The calculated extents of LM, CM, and L at an infinite time in the presence of mTG-23 are shown in **Table 1**. It is pertinent to mention that the inhibition of CM has been considered as the inhibition of fusion as the inner material (genetic material for viral fusion) cannot be transferred without proper pore opening.

**Figure 1.**
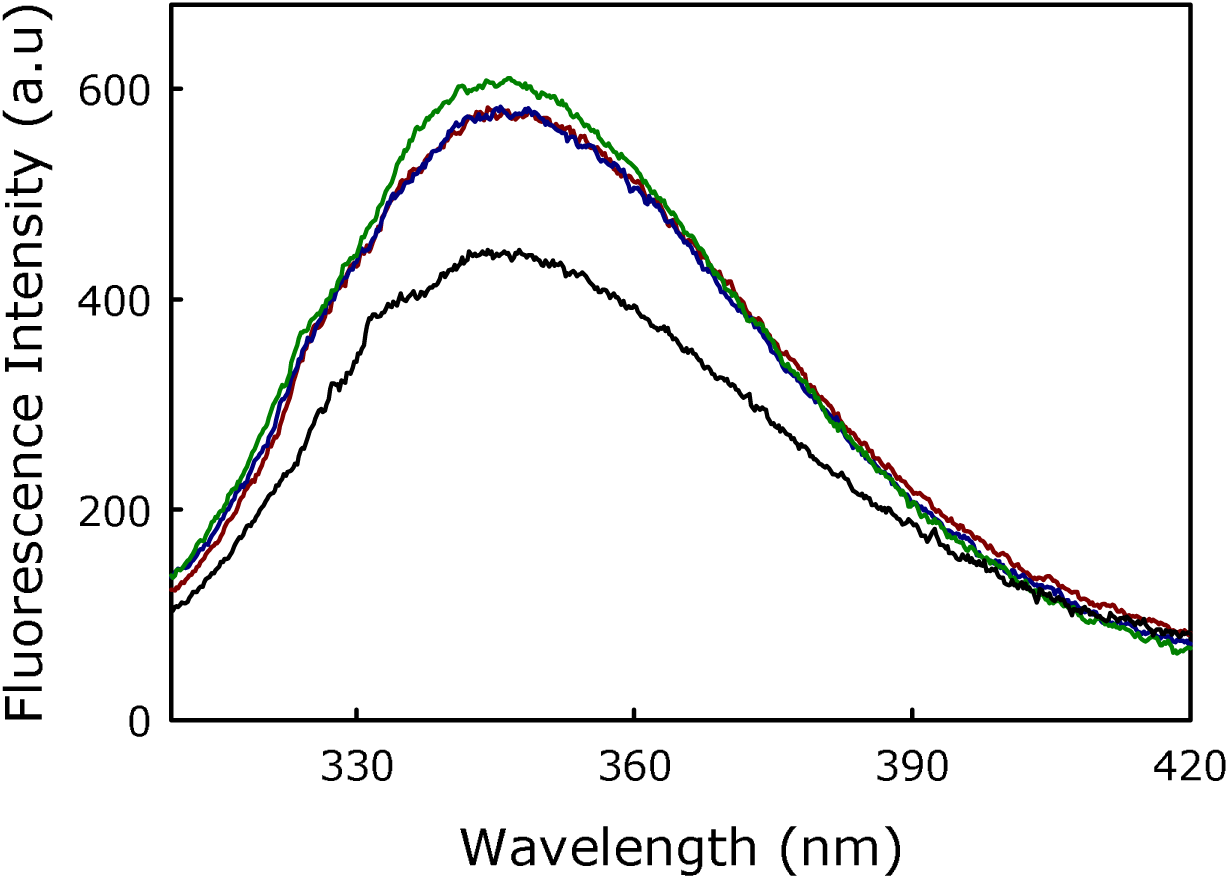
Fluorescence spectra of tryptophan present in mTG-23 in buffer (black), DOPC/DOPE/DOPG (60/30/10 mol%, dark red), DOPC/DOPE/DOPG/CH (50/30/10/10 mol%, blue) and DOPC/DOPE/DOPG/CH (40/30/10/20 mol%, green). Results are shown for a lipid-to-peptide ratio of 100:1 for measurements in the presence of membranes. Measurements were carried out in 10 mM TES, 100 mM NaCl, 1 mM CaCl_2,_ and 1 mM EDTA, pH 7.4 at a total lipid concentration of 200 μM. The data shown are the average of at least three independent measurements.

**Figure 2.**
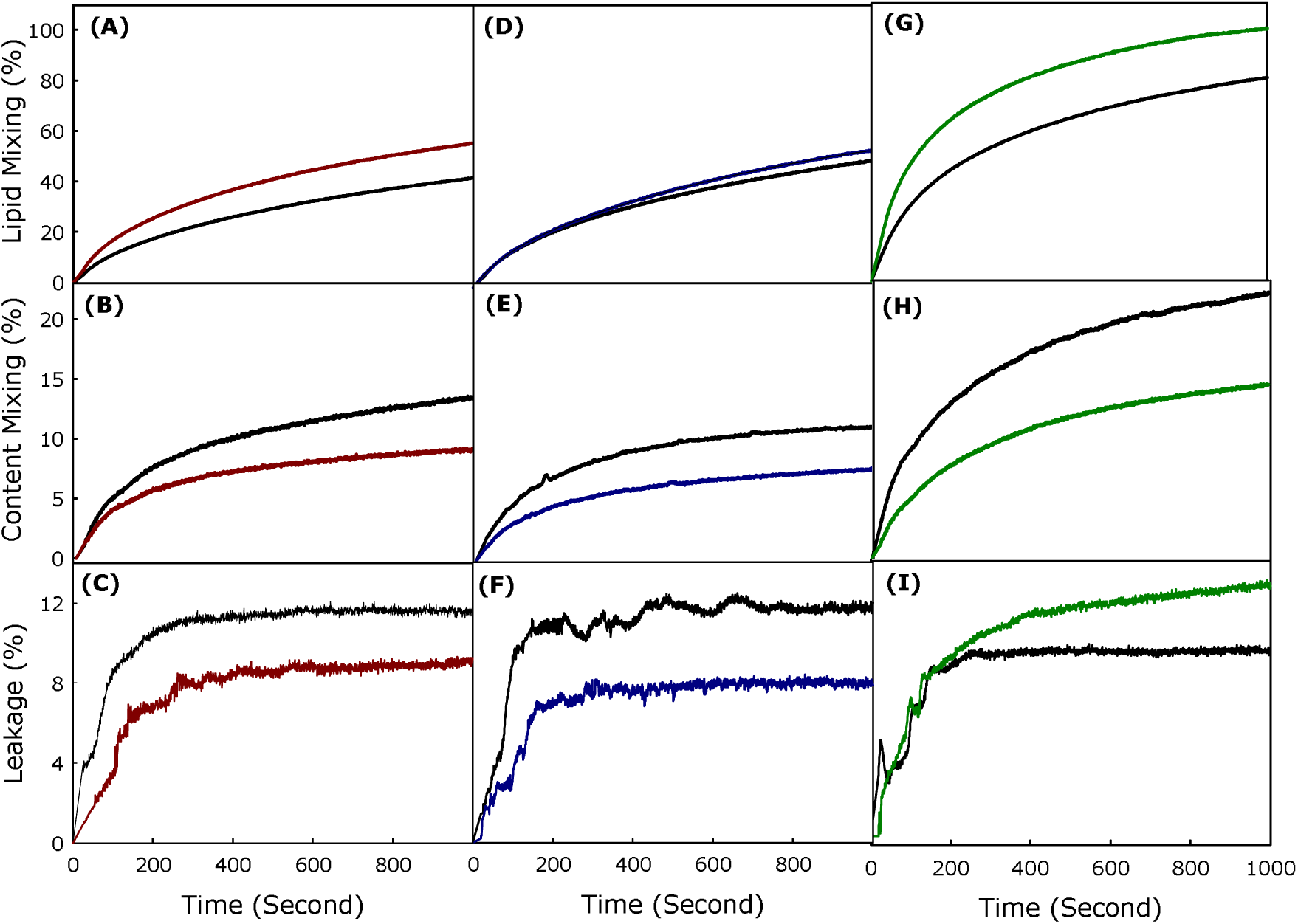
Plot of the effect of mTG-23 peptide on (A) lipid mixing, (B) content mixing, and (C) content leakage in DOPC/DOPE/DOPG (60/30/10 mol%), (dark red) and (D) lipid mixing, (E) content mixing, and (F) content leakage in DOPC/DOPE/DOPG/CH (50/30/10/10 mol%) (blue) and (G) lipid mixing, (H) content mixing and (I) content leakage in DOPC/DOPE/DOPG/CH (40/30/10/20 mol%) (green) SUVs. The black line in each figure represents the observable kinetics in the absence of peptide (control). Fusion was induced by 6% (w/v) PEG at 37 °C. Results are shown for a lipid-to-peptide ratio of 100:1. Measurements were carried out in 10 mM TES, 100 mM NaCl, 1 mM CaCl_2_, and 1 mM EDTA, pH 7.4 at a total lipid concentration of 200 µM. The data shown are the average of at least three independent measurements.

**Table 1.** The extent of lipid mixing, content mixing, and content leakage in the absence and presence of mTG-23 peptide in different lipid compositions. The data shown are mean ± SE of at least three independent measurements.

| Lipid Composition | Peptide | Lipid Mixing (%) | Content Mixing (%) | % of CM inhibition (Fusion Inhibition) | Content Leakage (%) |
| --- | --- | --- | --- | --- | --- |
| PC/PE/PG<br>(60/30/10) | Control | 57.4 ± 2.2 | 16.7 ± 0.6 | 41.0 | 11.7 ± 0.4 |
| | mTG-23 | 69.0 ± 2.7 | 9.8 ± 0.3 | | 8.7 ± 0.3 |
| PC/PE/PG/CH<br>(50/30/10/10) | Control | 67.8 ± 2.7 | 11.4 ± 0.4 | 29.0 | 13.6 ± 0.5 |
| | mTG-23 | 72.1 ± 2.8 | 8.1 ± 0.3 | | 8.3 ± 0.3 |
| PC/PE/PG/CH<br>(40/30/10/20) | Control | 92.7 ± 3.7 | 23.0 ± 0.9 | 33.0 | 11.1 ± 0.4 |
| | mTG-23 | 100 ± 3.9 | 15.5 ± 0.6 | | 14.4 ± 0.5 |

### Inhibitory effect of TG-23 versus mTG-23 peptide

Comparison of fusion inhibition by TG-23 and mTG-23 in membranes with different concentrations of membrane cholesterol brings out an important aspect of WD stretch in the inhibitory peptide (**Fig. 3**). It has been shown earlier that a WD dipeptide attached to a fatty acyl chain can inhibit membrane fusion.^41^ Interestingly, both tryptophan and aspartic acid have unique features to be located at the interfacial region of the membrane. The substitution of glutamic acid and alanine with tryptophan and aspartic acid near the C-terminus makes the peptide more versatile against the variation of lipid composition at least in terms of cholesterol content. The fusion inhibition has been reduced remarkably from 33% to 5% in the presence of TG-23 while the concentration of membrane cholesterol varied from 0 to 10 mol%, and it partially promotes fusion in the presence of 20 mol% membrane cholesterol. In contrast, mTG-23 inhibits fusion even in the presence of 20 mol% membrane cholesterol.

**Figure 3.**
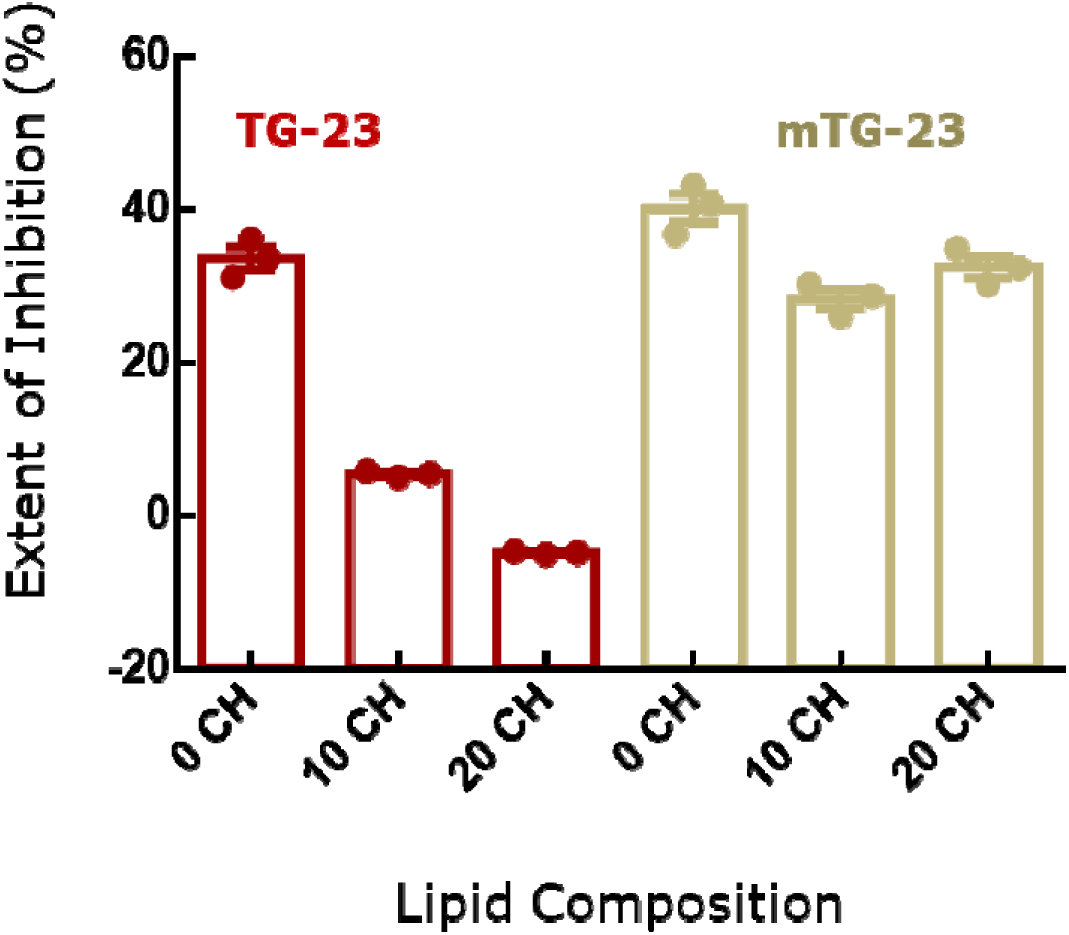
Comparison of percentage of fusion inhibition in terms of content mixing of TG-23 (dark red) and mTG-23 (yellow) peptides in DOPC/DOPE/DOPG (60/30/10 mol%), (0 CH) DOPC/DOPE/DOPG/CH (50/30/10/10 mol%) (10 CH) and DOPC/DOPE/DOPG/CH (40/30/10/20 mol %) (20 CH) SUVs. The data shown are mean ± SE of at least three independent measurements.

Therefore, placing WD in a strategic position significantly attenuates the inhibitory efficiency of the peptide. Importantly, WDs in the coronin 1 sequence are conserved across species,^24, 42^ and the protein plays a crucial role in avoiding lysosomal degradation of the live mycobacterium-loaded phagosome. Our results on mTG-23 support the importance of WD stretch in designing fusion inhibitors.

### Effect of mTG-23 peptide on the bilayer properties

To investigate the effect of the mTG-23 peptide on bilayer properties such as order and polarity at the interfacial region, the fluorescence properties of TMA-DPH were utilized. TMA-DPH is located at the interfacial region due to its polar trimethylammonium group, with an average distance of around 10.9 Å from the center of the bilayer.^43^ The fluorescence anisotropy of TMA-DPH provides information regarding the environmental rigidity^30, 44, 45^ and its fluorescence lifetime reveals the polarity of the interfacial region.^30, 44, 45^

### Effect of mTG-23 peptide on interfacial region

We have measured the peptide-induced change in fluorescence anisotropy of TMA-DPH in DOPC/DOPE/DOPG (60/30/10 mol%), DOPC/DOPE/DOPG/CH (50/30/10/10 mol %) and DOPC/DOPE/DOPG/CH (40/30/10/20 mol%) membranes in presence of varying concentrations of mTG-23 peptide. The fluorescence anisotropy results of TMA-DPH in different membranes have been studied and the results are shown in **Fig. 4A**. Our results indicate that mTG-23 disorders the interfacial region of both the non-cholesterol and cholesterol-containing membranes. The effect of the peptide is similar in the membrane containing 0 and 10 mol% of cholesterol; however, the change is less stiff in the presence of 20 mol% cholesterol-containing membranes. This could be due to the inherent rigidity of the membranes containing 20 mol% of cholesterol. Furthermore, the effect of mTG-23 peptide on the interfacial polarity was addressed by measuring the average fluorescence lifetime of TMA-DPH in DOPC/DOPE/DOPG (60/30/10 mol%), DOPC/DOPE/DOPG/CH (50/30/10/10 mol%) and DOPC/DOPE/DOPG/CH (40/30/10/20 mol%) membranes in presence of varying concentrations of mTG-23 peptide (**Fig. 4B**) to obtain information regarding the polarity of the environment surrounding the probe.^34, 46^ The presence of peptide partially reduces the interfacial polarity of the membranes and this is evident from the slight increase in TMA-DPH lifetime with increasing concentration of mTG-23. The partial increase in TMA-DPH fluorescence lifetime vis-à-vis reduction of interfacial polarity could be attributed to the peptide-induced exclusion of some water molecules from this region or maybe the lower polarity of the peptide compared to the lipid headgroups. To get a better estimate of the peptide-induced change in the membrane order we have calculated the apparent rotational correlation time of TMA-DPH using the Perrin equation (eqn. 7). **Fig. 4C** demonstrates the apparent rotational correlation time of TMA-DPH in different membranes in presence of varying concentrations of mTG-23. Like the fluorescence anisotropy, the apparent rotational correlation time drops significantly in the presence of the peptide irrespective of the lipid composition of the membrane. The enhancement of hemifusion formation (lipid mixing) in the presence of the mTG-23 peptide could be due to the peptide-induced disordering of the interfacial region of the membrane as lipids can undergo easy inter-bilayer movement in a disordered membrane. The peptide-induced reduction of interfacial polarity also supports its ability to increase hemifusion formation as membrane dehydration is often correlated to hemifusion formation.^47, 48^ However, this result does not explain the ability of the peptide to block pore formation.

**Figure 4.**
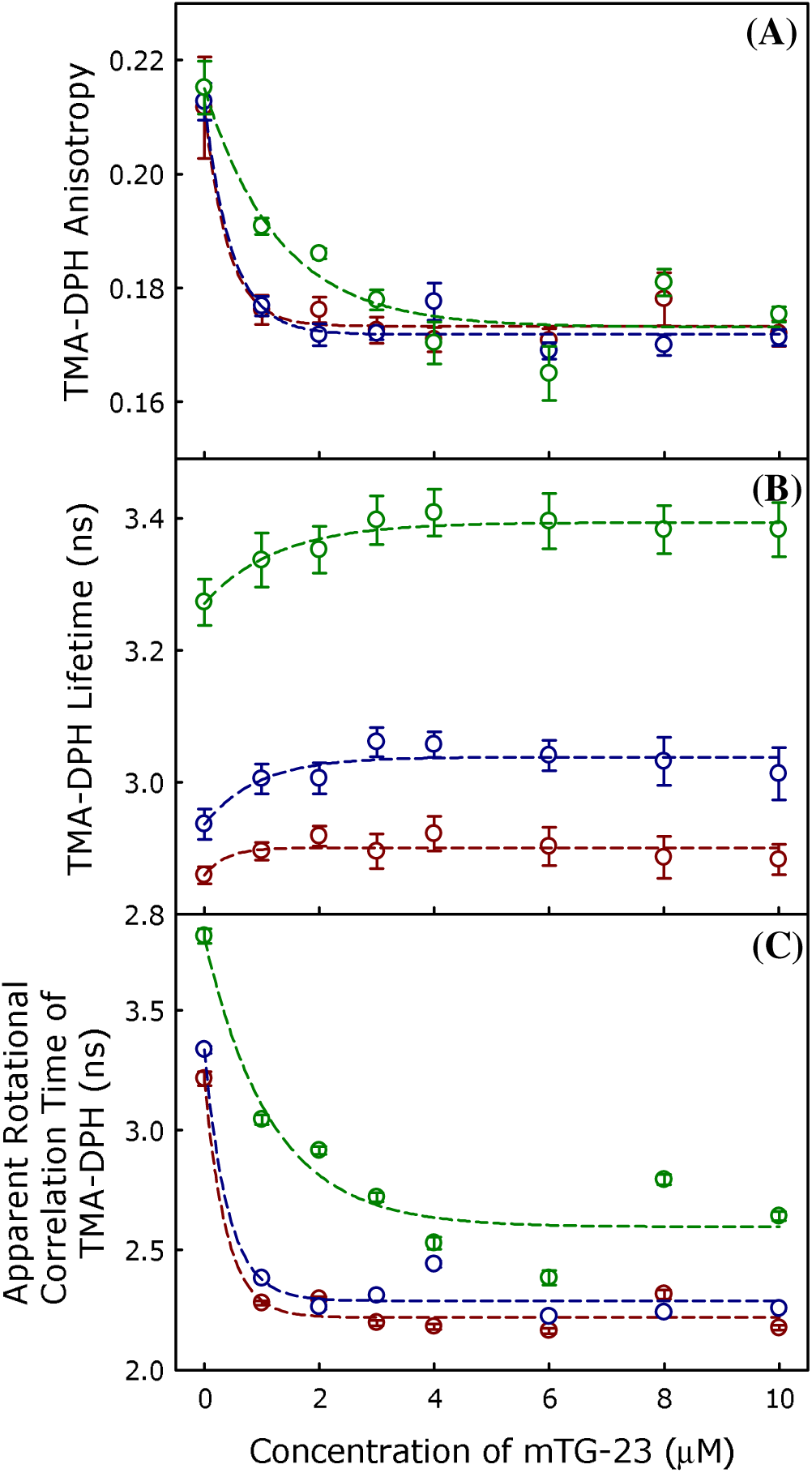
The plot of mTG-23 peptide-induced changes in the average (A) TMA-DPH anisotropy, (B) TMA-DPH fluorescence lifetime, (C) apparent rotational correlation time of TMA-DPH in DOPC/DOPE/DOPG (60/30/10 mol%) (dark red) DOPC/DOPE/DOPG/CH (50/30/10/10 mol %), (blue) and DOPC/DOPE/DOPG/CH (40/30/10/20 mol%), (green) SUVs as a function of peptide concentration. All the measurements were carried out in 10 mM TES, 100 mM NaCl, 1 mM CaCl_2_, and 1 mM EDTA, pH 7.4 at 37 °C, and a total lipid concentration of 200 µM. Data points shown are mean ± S.E. of at least three independent measurements.

### Antiviral effect of mTG-23 peptide

Enveloped viruses like HIV and Influenza transfer their genome to the host cells by fusing with either plasma or endosomal membranes of the host cell. We wanted to test whether mTG-23 inhibits virus-endosome / plasma membrane fusion. To test this, we evaluated the antiviral activity of mTG-23 against the Influenza A virus. To infect, the Influenza virus transfers its genome by fusion of endosomal membrane with viral envelope. We infected A549 and MDCK cells with X-31 (H3N2) influenza virus in the presence of varying amounts of the peptide. A549 is a human lung carcinoma cell line and is widely used as a preclinical model for Influenza infection. MDCK cell line is also routinely used for Influenza infection. The mTG-23 did not exhibit any significant cytotoxicity up to 50 µM as evaluated by MTT assay (**Supp. Fig. 3**).

**Fig. 5** shows that the influenza virus induces severe cytopathic effects (CPE) after 48 hours of infection. Interestingly, when the infection was carried out in the presence of increasing concentrations of the peptide, CPE was significantly reduced. To assess whether the reduction in CPE was associated with lower virus yield, media of the infected cells were collected after 48 hours of infection, and a hemagglutination assay was carried out. The hemagglutination assay provides a rapid readout of the relative viral load in terms of HA titer.^38^ The highest virus dilution that causes complete hemagglutination is considered the HA titration endpoint. HA titer is the reciprocal of the virus dilution in the last well with complete hemagglutination. **Fig. 5B** shows the reduction in HA titer in the presence of mTG-23; from 4 to 0, implying that mTG-23 inhibits influenza viral infection. To quantify the inhibitory effects, we evaluated the post-infection cell viability by MTT assay in the presence of varying amounts of mTG-23.

**Figure 5.**
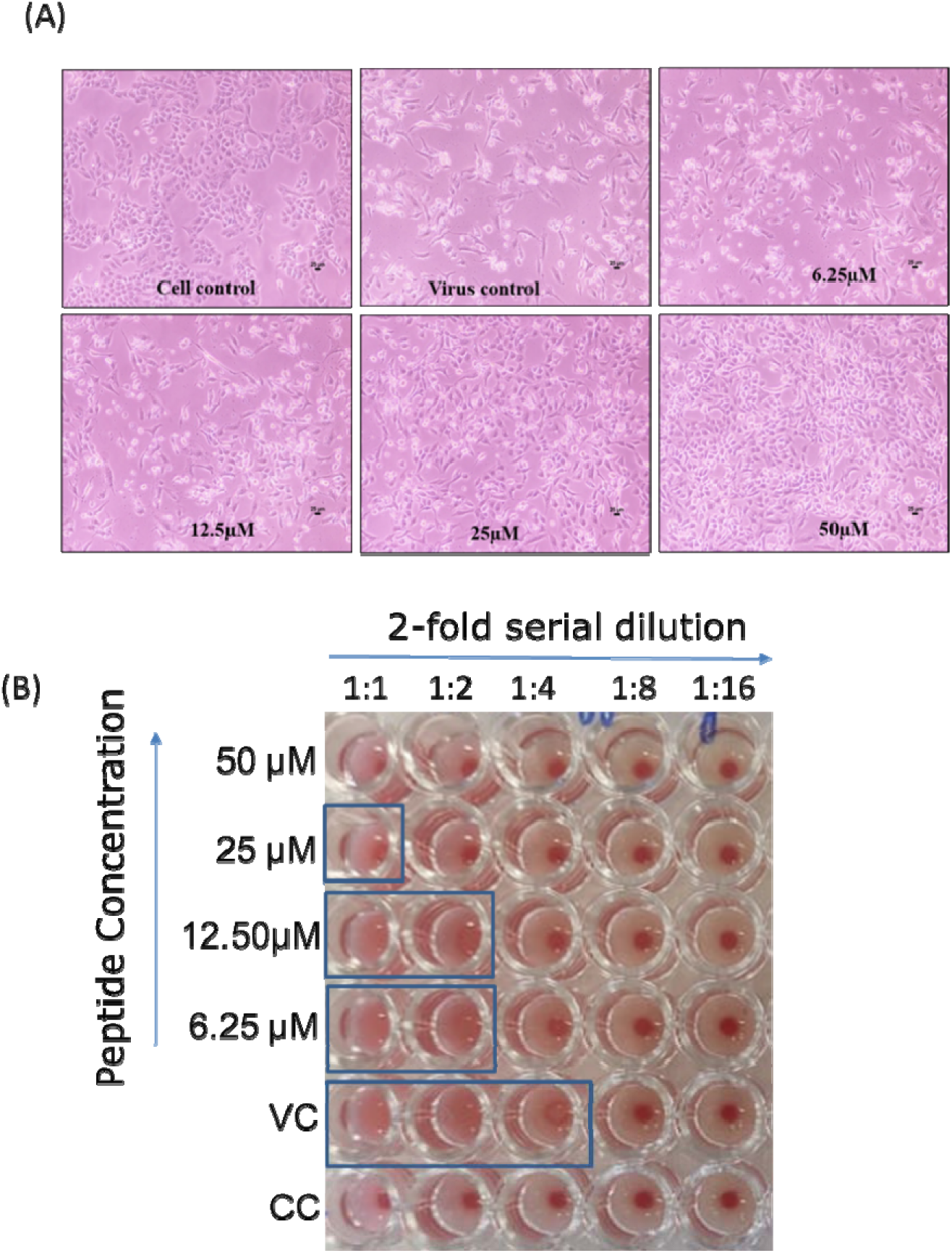
*In vitro* inhibitory activity of mTG-23 against Influenza A virus infection. (A) Light microscopy observation of a dose-dependent reduction in virus-induced cytopathic effect in the presence of mTG-23 in A549 cells after 48 hours of infection. A549 cells were infected with X31 (H3N2) influenza virus in the absence and presence of increasing concentrations of mTG-23. (B) Reduction in HA titer after 48 hours of infection in the presence of mTG-23. See the method section for details. VC and CC are samples from virus control and cell control, respectively.

**Fig. 6** shows a dose-dependent increase in cell viability (i.e., reduction in infection) of both A549 and MDCK cells. We have further quantified the post-infection cell viability as a normalized reduction in cytopathic effect (see methods for details). **Fig. 6C** and **D** show in A549 cells exhibits an EC_50_ of ∼ 20.45 ±1.60 µM while in MDCK cells it is ∼ 21.55 ±1.15 µM.

**Figure 6.**
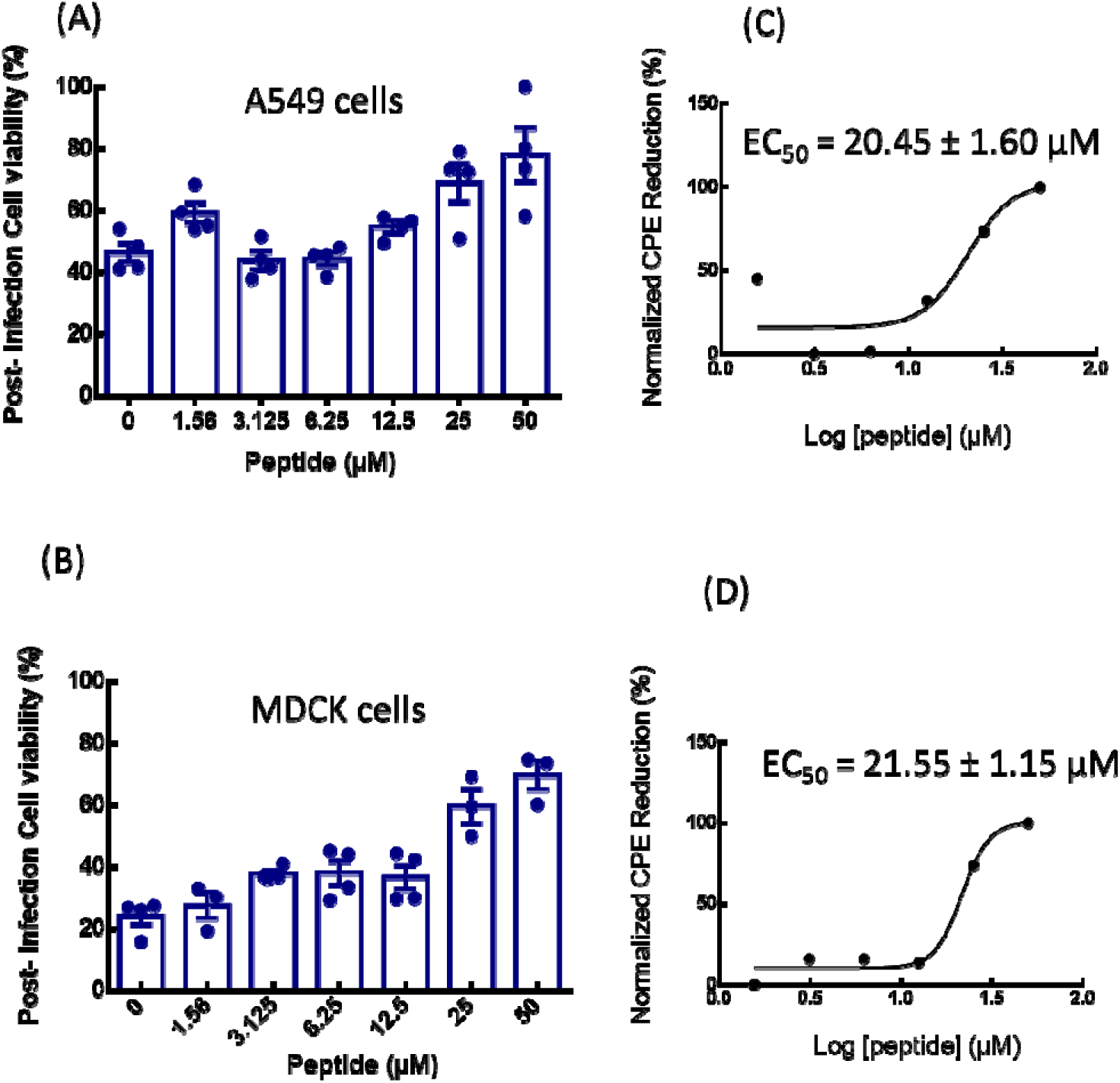
Quantification of *in vitro* anti-influenza activity of mTG-23. The effect of mTG-23 on the post-infection cell viability of A549 (A) and MDCK (B) cells after 48 hours of infection was determined by the MTT assay. Cell viability was normalized to control and represents mean ± SEM of at least 3 sets. Protection from CPE was normalized as described in the methods section and fitted to a dose-response curve. See the method section for details.

Next, we evaluated the expression level of M1 an Influenza viral protein in the absence and presence of mTG-23 by western blot. **Fig. 7**. shows a dose-dependent reduction in the level of M1 compared to the virus control in MDCK cells. A similar reduction was observed in mTG-23 against the X-31(H3N2) influenza A virus in A549 and MDCK cells. In the presence of mTG-23, there were no detectable changes in virus-RBC binding as detected by hemagglutination inhibition assay suggesting that mTG-23 did not affect virus binding to the host cells (data not shown). Based on our biophysical experiments with model membranes, our biochemical data suggests that mTG-23 inhibits influenza infection likely by preventing virus-endosome fusion. However, our data do not rule out any other mechanism behind the antiviral effect of mTG-23.

**Figure 7.**
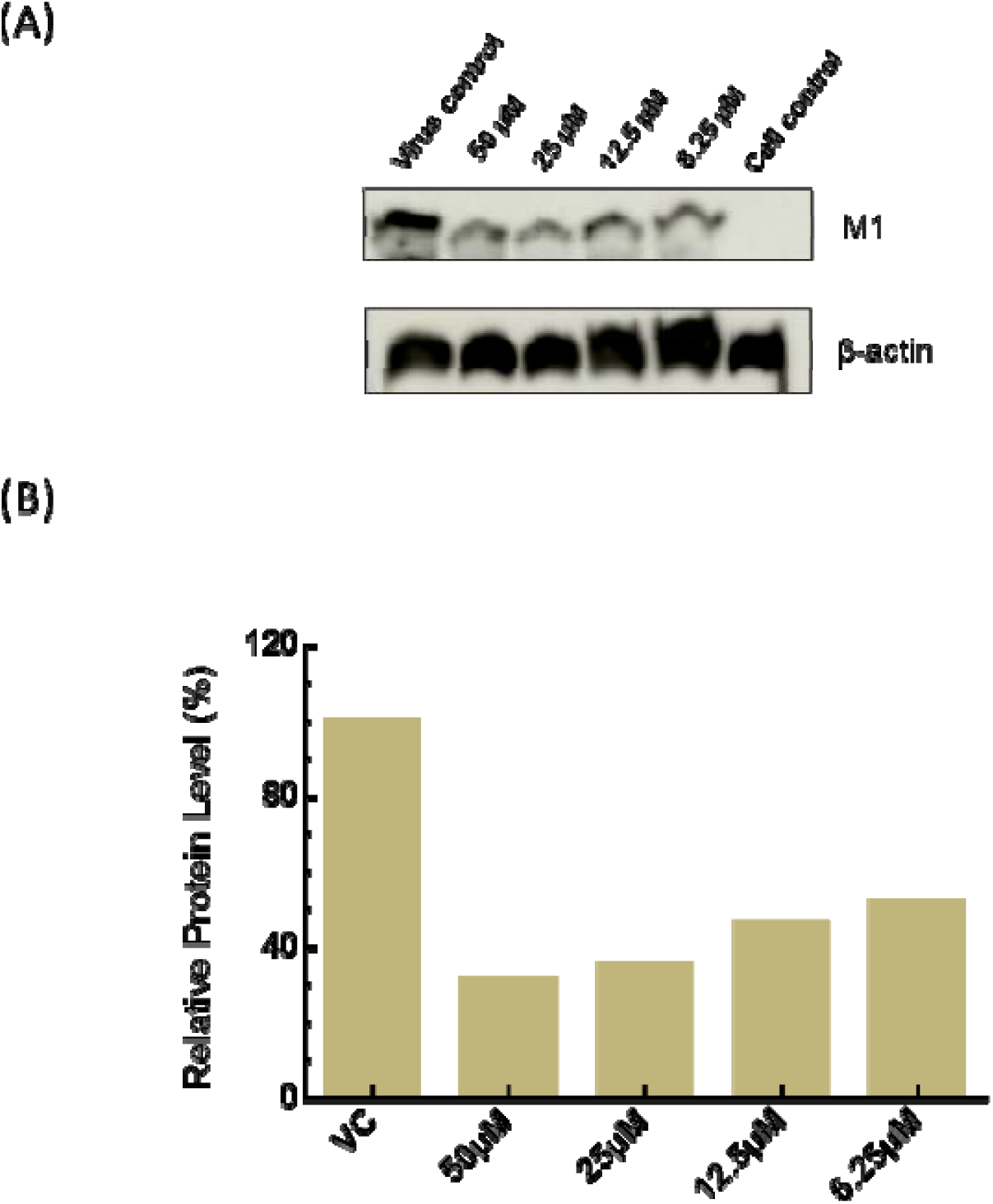
Dose-dependent reduction in viral protein expression. (A) A representative Western Blot showing the dose-dependent reduction in expression of Influenza M1 in MDCK cells after 48 hours of infection in the absence and presence of varying amounts of the peptide. β-actin was taken as a loading control. (B) Relative levels of M1 in the absence and presence of varying amounts of peptide.

## Discussion

The development of broad-spectrum anti-viral therapies is crucial to combat the continuously emerging and re-emerging viral diseases as the ‘one bug-one drug’ approach fails to be effective against new viruses. It is, however, difficult to develop broad-spectrum anti-viral agents that target the viral fusion proteins, as they change from virus to virus. Therefore, we have strategized to target the membrane and modulate its properties in such a way that the membrane would be less susceptible to fusion. We have earlier shown that Coronin 1-derived WD-containing peptides can block PEG-induced fusion of SUVs by modulating membrane physical properties.^22, 49^ The presence of WD helps the peptide to modulate the interfacial properties of the membrane as they partition at the membrane interfacial region because of their mixed polarity. In this work, we have modified a peptide sequence derived from Coronin 1 by introducing another WD repeat near the C-terminus and developed a new peptide called mTG-23. The mTG-23 contains two WD repeats at two termini of the peptide, therefore, it is expected that the peptide would be locked at the membrane interface through terminal WD repeats as both tryptophan and aspartic acid prefer interfacial localization.^50^ The middle part of the peptide is expected to be partitioned into the membrane because of the chain hydrophobicity. Overall, the peptide is expected to behave as the miniature version of the coronin 1 protein which is known to coat the phagosome and inhibits membrane fusion.^23^ As expected, the peptide is indeed preventing model membranes from fusing. We hypothesize that mTG-23 may also inhibit viral fusion and consequently prevent viral infections. We have tested this hypothesis with the Influenza A virus (IAV). After the IAV particles bind to host cells, they are taken in through endocytosis. While being transported to the perinuclear region, the viral particles encounter low pH levels within the late endosomes, which causes the viral surface glycoprotein hemagglutinin (HA) to undergo a series of conformational changes, exposing the fusion peptide. The fusion peptide connects the viral envelope with the endosomal membrane and is responsible for the fusion of the viral envelope with endosomal membranes, allowing the transfer of viral RNA into the cytoplasm.^7, 51^ Our results show that mTG-23 inhibits the Influenza A virus infection in A549 and MDCK cells. Since we co-treated cells with the virus and the peptide, mTG-23 likely inhibits the early steps in the infection cycle: binding, entry, and fusion.^52^ Since we did not observe any effect on the binding of the virus to RBC by hemagglutination inhibition assay, this suggests that the virus binding to the host cells is not affected by mTG-23. Therefore, it is plausible that mTG-23 is inhibiting the fusion of the viral envelope and endosomal membranes. However, our phenotypic assays do not exclude the possibility that mTG-23 might affect other steps in the infection cycle. A detailed mechanistic study to decipher the mechanism of the antiviral effect of mTG-23 will be carried out next. Nonetheless, our results surmise that mTG-23 might prevent infection of other viruses which involve membrane fusion as a crucial step in the infection cycle. Because of the antigenic shift and antigenic drift, Influenza, and other viruses constantly evolve and evade our immune system, gaining resistance to existing antiviral drugs and reducing vaccine effectiveness. Consequently, novel sustainable antiviral strategies against mutant strains of deadly viruses like influenza are needed continuously to avoid the next pandemic.^53^ Rationally designed peptides such as mTG-23 that target the fusion pathway represent a novel pan-antiviral strategy. Moreover, our results unfold a new avenue to inhibit IAV infection. Importantly, there are not ample therapeutic interventions available to combat IAV infection^54^ and our approach might help to prevent one of the most common but contagious viral infections.

### Author contributions

M.S.P and G.P.P carried out the spectroscopic and model membrane experiments and wrote the initial draft manuscript, B.Q and V.V performed the biochemical and virology experiments and wrote the initial draft manuscript, S.H, and H.C conceptualized and supervised the work, acquired funding, and wrote and edited the final manuscript.

## Acknowledgments

This work was supported by the Core Research Grant (CRG/2021/001515) of the Science and Engineering Research Board (SERB), the SERB-Science and Technology Award (STR/2021/000029) for Research, New Delhi to HC, and Ramalingaswami Re-entry Fellowship (BT/RLF/Re-entry/11/2020) to S.H. H.C. and M.S.P. thank the University Grants Commission for the UGC-Assistant Professor position, and SERB for fellowship, respectively. BQ thanks CSIR for the award of Junior Research Fellowship. S.H. thanks the Division of Virus Research and Therapeutics, CSIR-CDRI for providing the infrastructure. X-31 Influenza A virus was a kind gift from Dr. Joshua Zimmerberg from NIH, Bethesda. We thank DST, New Delhi, and UGC for providing the instrument facility to the School of Chemistry, Sambalpur University under the FIST and DRS programs, respectively. We thank members of the Chakraborty and Haldar laboratories for their comments and discussions.

