## Supplemental figures for "Developing peptide-based fusion inhibitors as an antiviral strategy utilizing Coronin 1 as a template"

#### Electronic supplementary information (ESI)

### Supplementary Figures

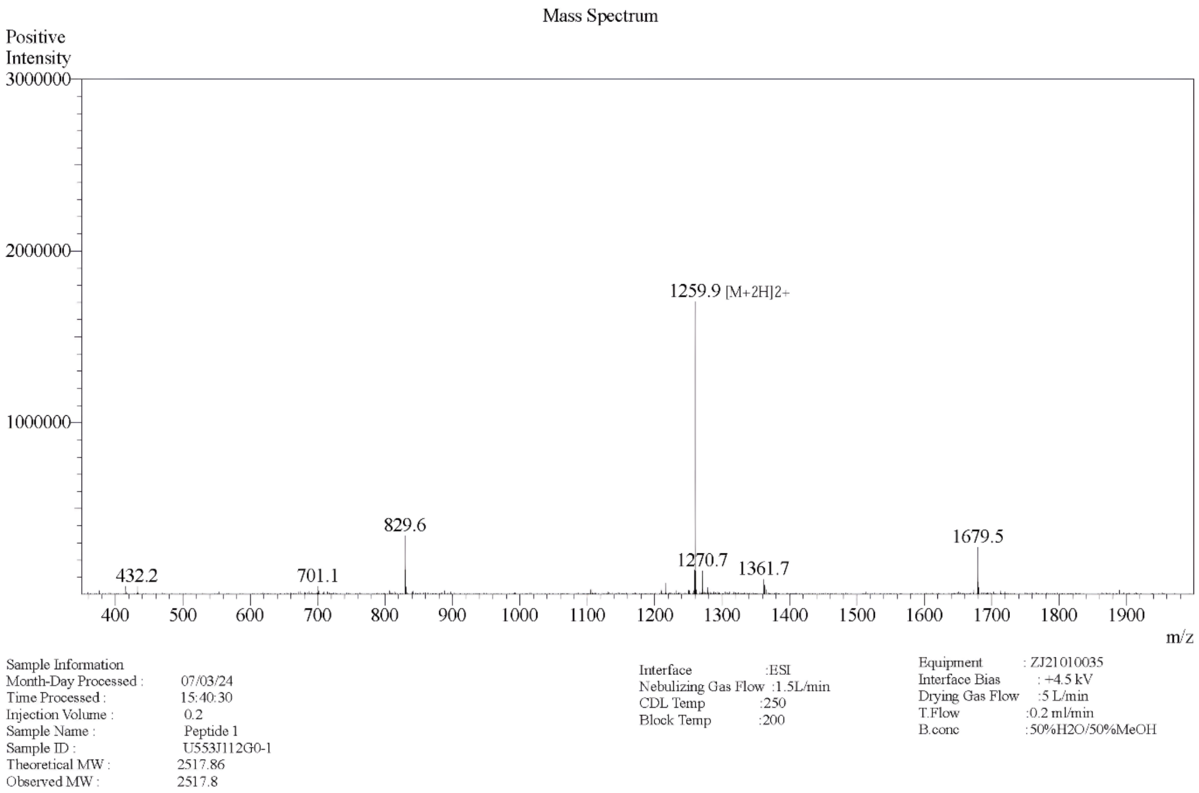


**Supp. Fig. 1**. Mass spectrum of mTG-23 peptide.

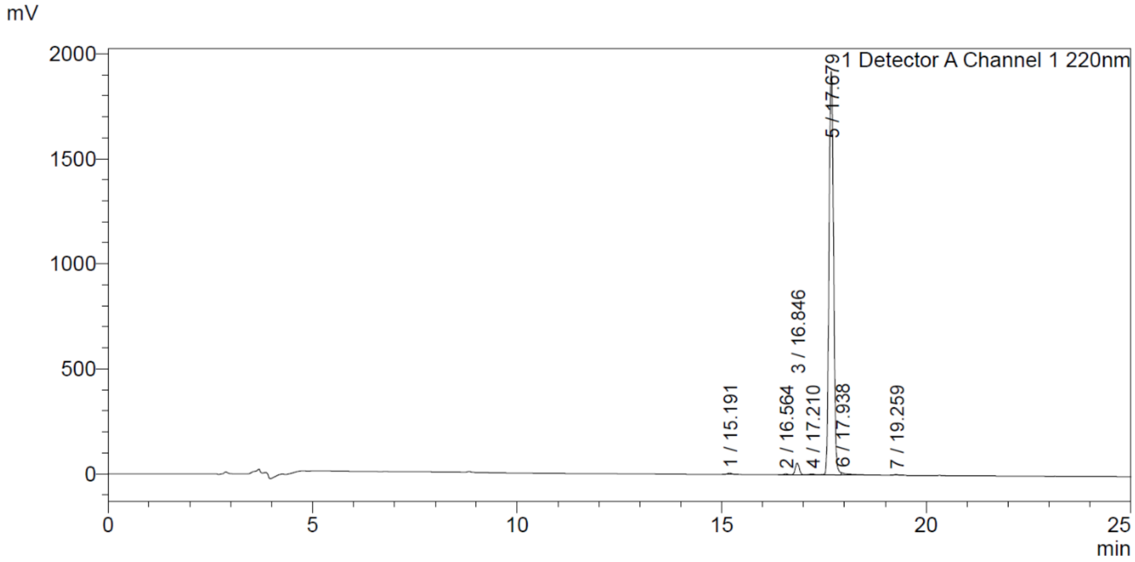


**Supp. Fig. 2**. HPLC profile of mTG-23 peptide. The peptide has been eluted through Inertsil ODS-SP column using 0.05% trifluoroacetic in 100% acetonitrile (v/v) as a solvent in gradient mode at 1 ml/min flow rate.

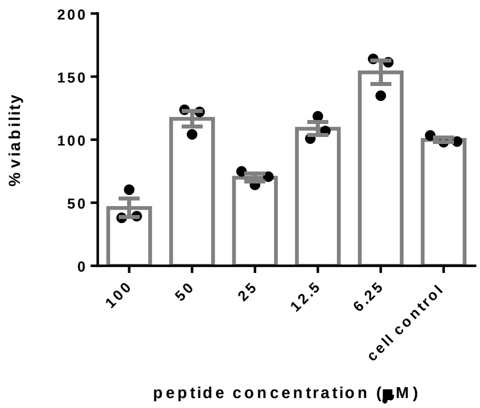


**Supp. Fig. 3**. A549 Cell viability in the presence of mTG 23. Data represent mean ± SEM of at least 3 sets.

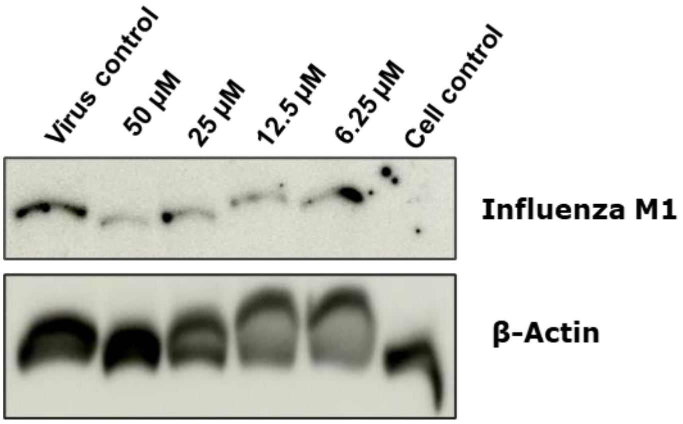


**Supp. Fig. 4.** A representative Western Blot showing the dose-dependent reduction in expression of Influenza M1 in MDCK cells after 48 hours of infection in the absence and presence of varying amounts of peptide. β-actin was taken as a loading control.
